# Variability of transcriptional response to water deficit and low temperature in leaves of wheat *Triticum aestivum* L. of extensive and intensive type

**DOI:** 10.64898/2026.03.16.711993

**Authors:** I.V. Gorbenko, Yu.M. Konstantinov, S.V. Osipova

## Abstract

The paper presents the results of a comparative analysis of gene networks activated by water stress and low temperatures in extensive (Saratovskaya 29, S29) and intensive (Yanetskis Probat, YP) wheat varieties during the seedling development stage. It is concluded that the creation of the S29 variety, which occurred through complex stepwise hybridization and selection for morphological traits, productivity, and grain quality traits, resulted in the emergence and inheritance of regulatory gene networks involving proteins with the CC domain, as well as the BTB/POZ and TAZ domains, which have an increased affinity for transcription factors involved in the transcriptomic response to changing external conditions. It was established that, at the transcriptomic level, the S29 variety is characterized by a transition to an energy saving mode to maintain the activity of the Calvin-Benson cycle under the water deficit conditions and the inhibition of proteolytic processes at low temperatures. The transcriptional response of the high-yielding YP variety to 24-hour low-temperature treatment was more active and involved a larger number of genes compared to the S29 variety. Identifying varietal variability in molecular genetic mechanisms of resistance to abiotic stressors facilitates the development of marker-assisted and genomic selection technologies for common wheat.

**Key message:** The extensive S29 variety was characterized by its transition to energy-saving mode to maintain the Calvin-Benson cycle under water deficit and a reduction in proteolytic processes under low temperature.

## Introduction

Advances in RNA sequencing (RNAseq) technologies and transcriptome analysis methods facilitate the deciphering of gene networks activation under stress conditions, both in model plants with small genomes and in hexaploid wheat *Triticum aestivum* L. (2*n* = 6x = 42, BBAADD), with 16 Gbp size genom (The et al., 2018). RNAseq has become a key tool for investigation of wheat responses to fluctuations in environmental conditions, allowing us to detect both general and specific, even minor, changes in gene expression and identify genes involved in the formation of resistance to certain unfavorable environmental factors (Andleeb et al., 2024). The latter is crucial for the creation of new varieties of this cereal, which is crucial for solving the world’s food problem, using marker-assisted and genomic selection technologies.

Leaves are the main organ of plants that perform photosynthesis and an important organ that perceives environmental signals for adaptation (Biswal & Kohli, 2013). Using RNAseq of flag leaves and roots Wang et al. (Wang et al., 2020) discovered genes modulated by iron deficiency stress and demonstrated the role of several iron ligands, iron transporters, and regulatory mechanisms involved in iron homeostasis in wheat plants. Based on transcriptomic data analysis of wheat leaves subjected to salt stress, Amirbakhtiar et al. (Amirbakhtiar et al., 2021) and Prasad et al. (Prasad et al., 2022) revealed variety-specific features of molecular reactions of varieties with different salt sensitivity and suggested genes for improvement of wheat salt-resistance. Gálvez et al. (Gálvez et al., 2019) successfully used RNAseq of wheat leaves in studies of the genomic architecture of drought responses, discovering compact clusters of genes that were differentially expressed in wheat cultivar Chinese Spring under drought stress in the field. Transcriptome analysis of leaf tissue showed that two drought-tolerant bread wheat varieties, TAM 111 and TAM 112, well adapted to the conditions of the same region of North America and related by approximately 40% common ancestry, implemented different mechanisms of drought tolerance at the molecular level (Chu et al., 2021). At the transcriptome level, the TAM 112 variety responded more actively to water stress than the TAM 111 variety. TAM 112 demonstrated more active regulation of the expression of genes involved in protein phosphorylation processes, genes with adenyl ribonucleotide binding function, and genes involved in transcription regulation and transmembrane transport. The authors suggest that the reduced expression of these genes in TAM111 helped conserve energy for carbohydrate transport from stems and ears to caryopses or even led to adjustments in gene expression to interrupt seed development in some ears to preserve overall yield. At the phenotypic level, differences in gene expression manifested themselves in the formation of a higher number of sterile spikelets per ear in TAM 111 and higher yield during prolonged drought in TAM 112. In the same study (Chu et al., 2021), the importance of genes encoding proteins of intracellular membrane organelles (chloroplasts, mitochondria, and endoplasmic reticulum) in response to water stress was noted. Transcriptome analyses of flag leaves of different wheat varieties under changing environmental conditions revealed intraspecific variability in stress responses at the molecular level. This variability was explained both by differences in plant growing conditions across experiments (Gálvez et al., 2019) and by intraspecific genomic variability. After conducting whole-genome sequencing and analysis of genomes of 15 varieties, Walkowiak et al. (Walkowiak et al., 2020) concluded that intraspecific variability in common wheat genomes is the result of breeding efforts aimed to improve adaptation to various environmental conditions, stress tolerance, yield, and grain quality.

At early spring in Eastern and Western Siberia spring wheat plants are at the seedling growth and early tillering stages, and often are subject to droughts with recurrent temperature drops. Successful wheat adaptation depends on particular conditions of this phase, as primary roots, productive shoots, internodes, and stem leaves develop during this period (Efremova & Chumanova, 2023). Given the key role of early stages of wheat plant development, Konstantinov et al. (Konstantinov et al., 2021) conducted whole-genome sequencing of third leaf tissue from the common wheat varieties Saratovskaya 29 (S29) and Yaneckis Probat (YP) at the fourth leaf stage under optimal conditions (Control), under water deficit (WD), and under low positive temperatures (+4°C) for 6 (6H) and 24 (24H) hours. Konstantinov et al. (Konstantinov et al., 2021) identified common adaptation strategies to abiotic stress factors for S29 and YP. The expression of 43 genes changed similarly in response to water deficit and low-temperature stress. The expression consistency between homeoallelic gene sets and the distribution of SNPs across the genome were also analyzed. The authors compared differentially expressed genes (DEGs) for the S29 and YP varieties with known quantitative trait loci (QTLs) associated with antioxidant enzyme activity under water stress. However, the S29 and YP varieties showed a relatively low overlap of DEGs, less than 10% for each stress type. This may be due to significant differences in gene expression in response to the studied stresses and to different molecular strategies for adaptation to adverse conditions in these varieties. In S29 wheat, a drought-tolerant, extensive-type variety, the expression of AAA family ATPases, signaling and regulatory proteins with an F-box domain, and alpha/beta hydrolase superfamily proteins increased under water stress. The intensive wheat variety YP was characterized by reduced expression of transmembrane amino acid transporters. However, given the continuous advances in comparative transcriptomics approaches and methods, there is a need to further research to identify the potential diversity of molecular strategies for wheat adaptation to abiotic stress factors.

The aim of the current work was to identify differences in the molecular adaptation strategies of these spring hexaploid wheat varieties, but with different pedigrees and different resistance to unfavorable external factors at the seedling growth stage using the gene network method.

## Materials and methods

For the analysis we took relative expression values of 137,000 genes obtained for the common wheat varieties S29 and YP grown under optimal controlled conditions (S29.control, YP.control), under water deficit (S29.WD, YP.WD), and at 4°C for 6 h (S29.6H, YP.6H) and 24 h (S29.24H, YP.24H). The relative gene expression values were calculated using the formula ((x / sum_red) * 1,000,000, where x is the number of reads mapped to the gene; sum_red is the total number of mapped reads in the library) are published in Supplementary Data S4 of Konstantinov et al. (Konstantinov et al., 2021).

The data analysis was conducted in R programming language environment (R Development Core Team, 2022). Co-expression analysis was performed using WGCNA-based R package CEMiTool (Russo et al., 2018). For particular correlation expression analysis, we took only the strong correlations (R_Pearson_ > 0.8). The enrichment analysis was performed using ClusterProfiler R package (Yu et al., 2012). Heatmaps were made using ComplexHeatmap R package (Gu et al., 2016), other graphics were made using ggplot2 (Wickham, 2016) and ggpubr (Kassambara, 2026). Networks construction and analyses were performed using igraph R package (Csárdi et al., 2026), plots were created using the ggraph R package (Pedersen, 2025).

## Results

### Gene co-expression modules in S29 and YP varieties

Analysis of expression values of 137.000 of genes of wheat varieties S29 and YP revealed probable co-expression for 510 genes which were organized in 6 co-expression modules (CEMs). The full genelist of CEMs is available in Supplementary Data S1. The result of enrichment analysis of CEMs is shown in Fig. 1.

**Figure 1.**
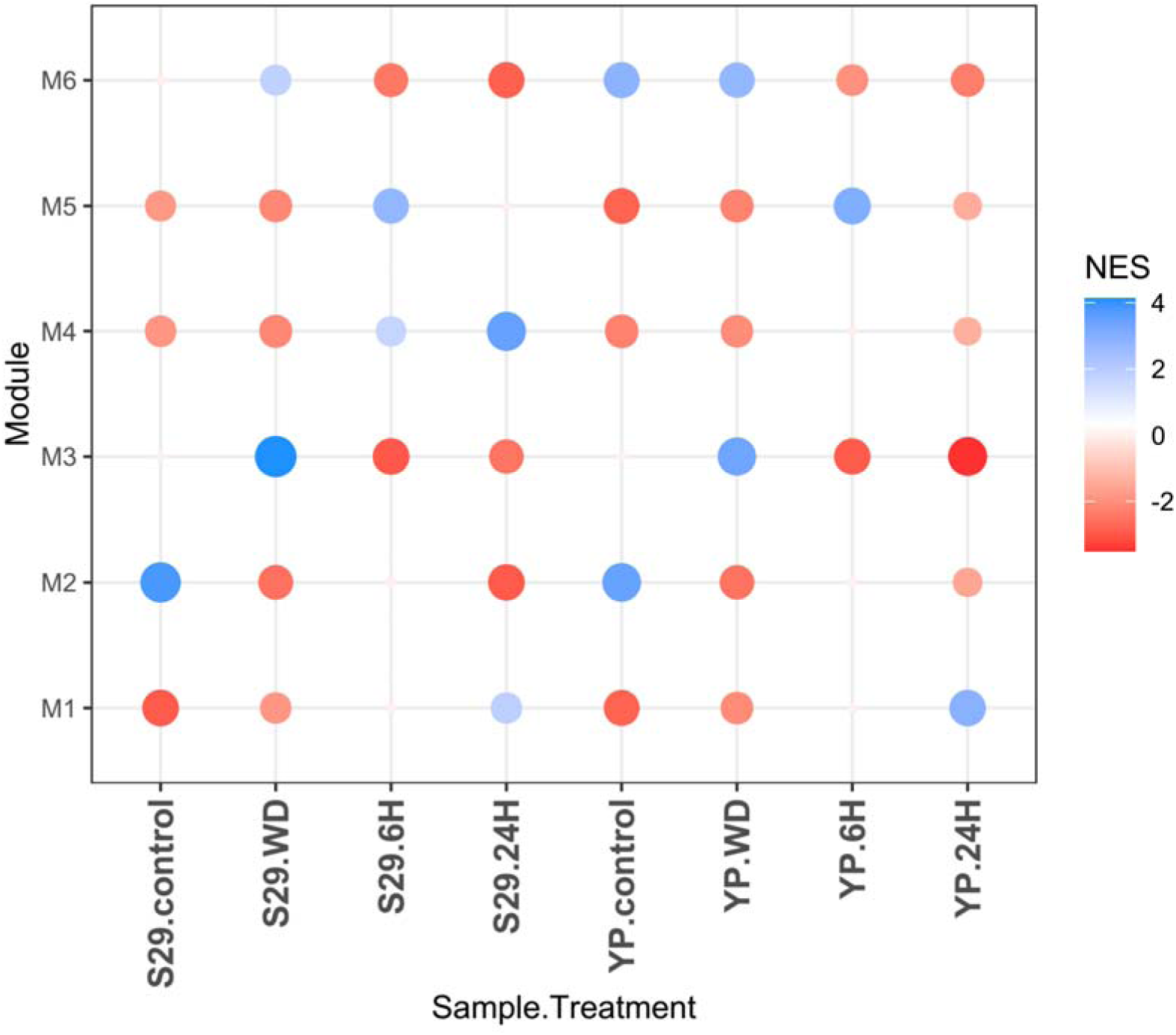
The enrichment of co-expression modules in S29 and YP in control, under the water deficit (WD), and under the low positive temperature (+4L) treatment for 6 hours (6H) and 24 hours (24H). NES stands for Normalized Enrichment Score and was calculated as z-scaled relative expression of CEM genes in a sample.

Module M1 was consisted of 149 genes (Supplementary Data S1). According to the overrepresentation analysis of Gene Ontology terms, the genes of this module are in general associated with stress-reactions (Fig. 2). The genes of M1 were located in all 42 chromosomes of bread wheat. In descendance In order of decreasing number of M1 genes, homeologous groups of chromosomes were distributed as follows: 5 (32 genes) > 2 (26) > 4 (22) > 7 (20) > 3 (19) = 6 (19) > 1 (11).

**Figure 2.**
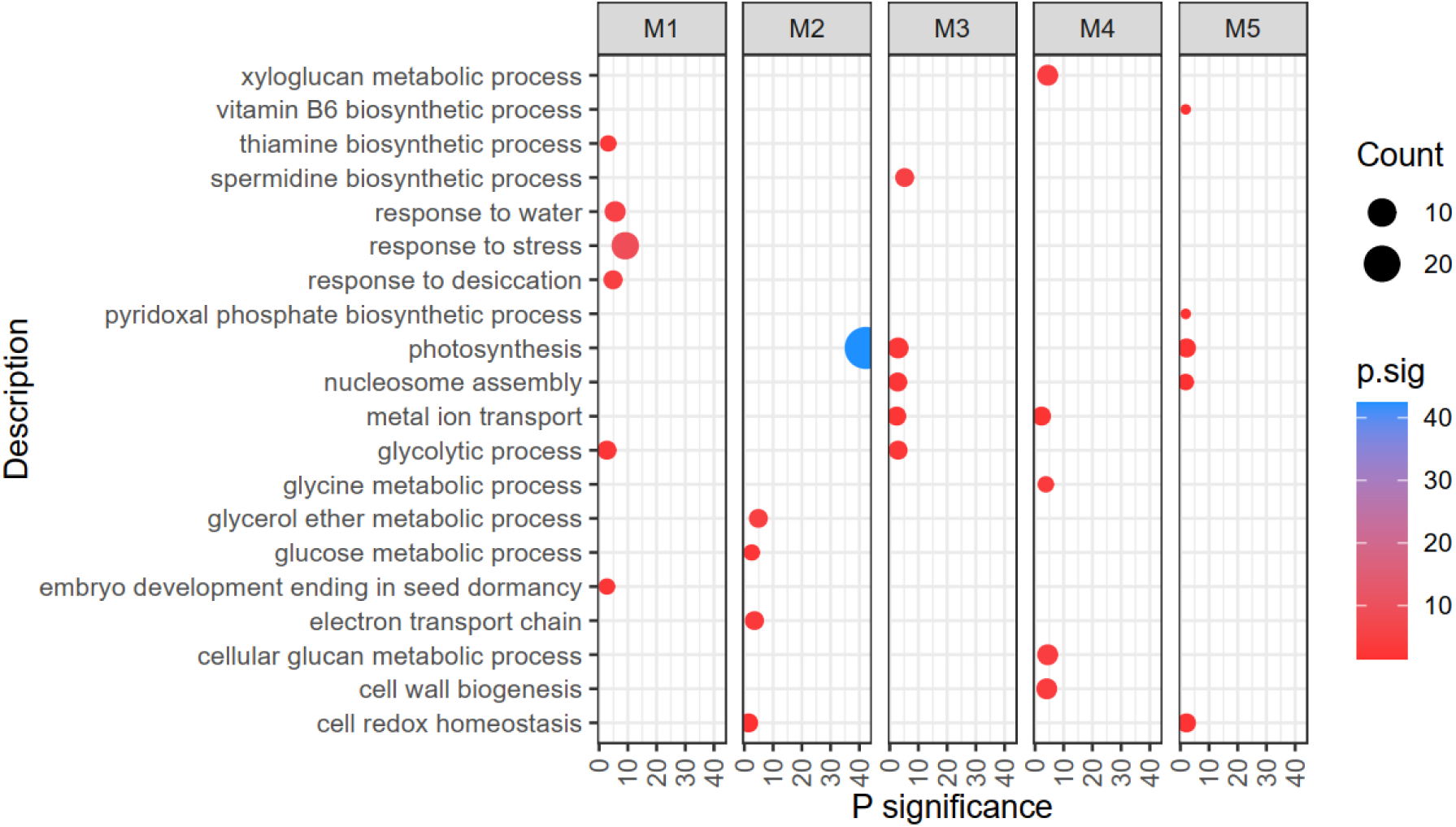
The results of overrepresentation analysis of GO terms in 5 co-expression modules (top-5 terms are shown). The *P* significance, used here as overrepresentation score, was calculated as (-1)*log(*P_BH_*), where *P_BH_*is *P-*value adjusted via the Benjhamini-Hochberg method.

The expression of M1 genes was activated to the same extent in both varieties under the WD and 6H conditions. Under the 24H conditions the enrichment of M1 was noticeably stronger in YP then in S29 (Fig. 1). The hub-genes (genes with maximum adjacency in co-expression module network) of M1 consisted 3 homeologs encoding Actin depolymerizing factor; a dehydrin protein which closest known ortholog is a cold-induced DHN5 of *Hordeum vulgare (Q9SPA7_HORV*); 2 homeologs of late embryogenesis abundant protein (LEA); senescence-associated Zink-finger type transcription factor (TF) which closest known ortholog is FLZ of *A. thaliana* (*AT4G17670.1*); WPI6 protein induced in response to low temperatures and salt stress. The expression of 7 of the top-10 hubs of module M1 was increased in S29.6H, and in S29.24H only 3 of them. On the contrary, in YP.6H the expression of 3 of these genes was increased, and in YP.24H – the expression of all 10 of the mentioned was increased (Fig. 3). Under the WD conditions the expression of M1 hub genes was unaffected (Fig. 3). The centrality degree of M1 hub genes was noticeable high (∼300, Supplementary Data S2).

**Figure 3.**
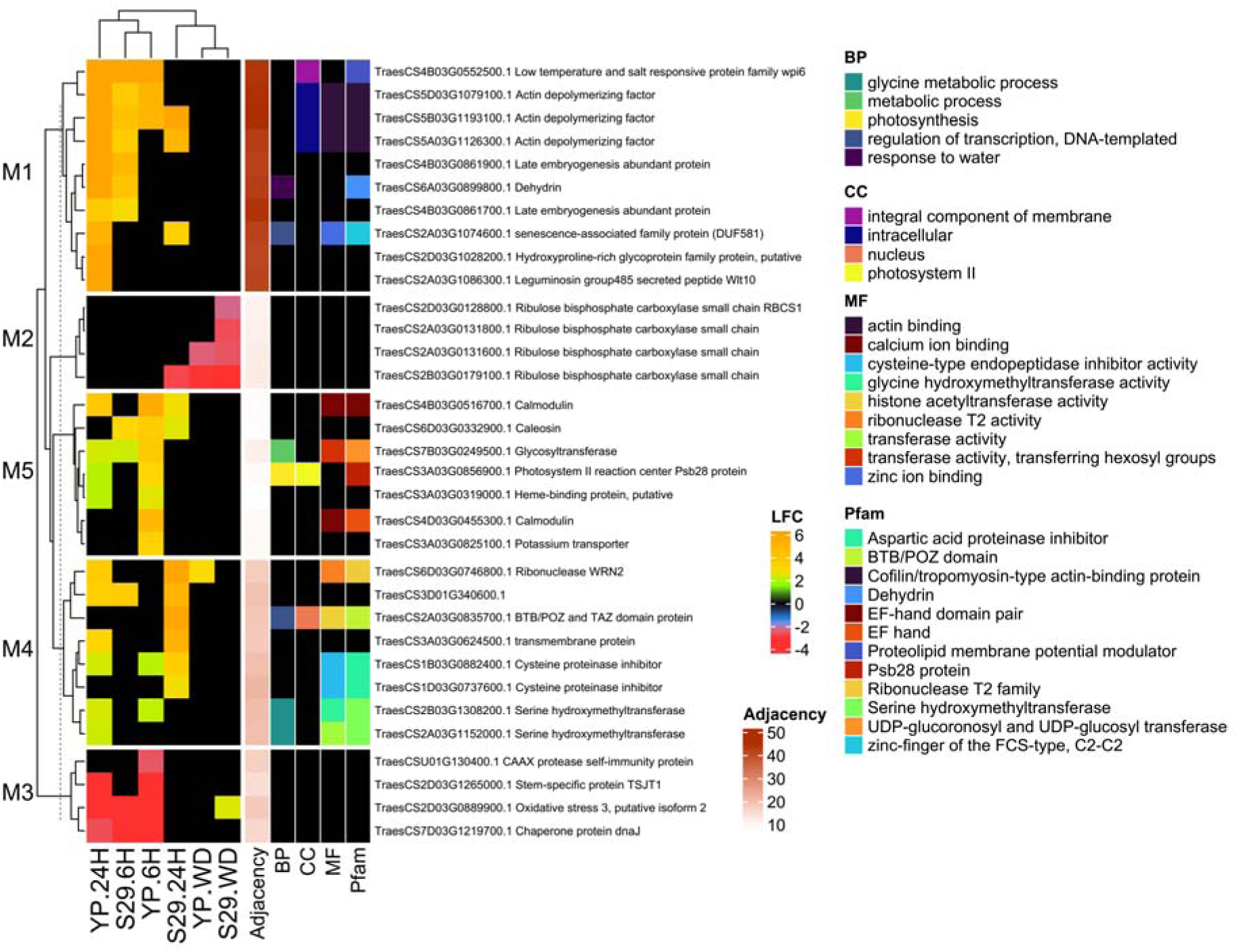
The expression of CEM hub genes (nodes of maximal adjacency). LFC stands for Log Fold Change. Additional color annotation provides hubs features: Adjacency, BP – main GO Biological Processes term, MF – main GO Cellular Components term, MF – main GO Molecular Functions term, Pfam protein family.

Module 2 (M2) consisted 103 genes (Supplementary Data S1). Homeologous groups of chromosomes were distributed as follows: 2 (34 genes) > 5 (23) > 7 (16) > 6 (13) > 3 (8) > 1 (6) > 4 (3) in decreasing order of M2 genes number. According to GO terms, M2 is dominated by genes associated with photosynthesis processes and contains genes regulating cellular redox homeostasis (Fig. 2). Under the WD, M2 was suppressed in both varieties approximately equally, whereas under the 24H M2 was suppressed to a greater extent in S29. Expression of the M2 hub genes encoding the small subunit of Rubisco decreased under the WD in both varieties, with the expression of all four homeologs decreased in S29 and only two of them in YP (Fig. 3). In addition to the small subunit of Rubisco, the PsaG and PsaK subunits of the photosystem I (PSI) reaction center, the PSII-T subunit, and the PsaH subunit of PSI were among the hub genes of M2. The proteins with maximal adjacency in PPI network of M2 had a relatively low degree of centrality (12, Supplementary Data S2).

Module 3 (M3) consisted of 97 genes (Supplementary Data S1). In decreasing order of the number of M3 genes, homeologous groups of chromosomes were distributed as follows: 2 (23 genes) > 3 (15) = 4 (15) = 6 (15) > 7 (10) > 5 (10) > 1 (8). M3 was activated in response to the WD, to the greater extent in the S29 variety, and was suppressed under the low temperature treatment, more strongly in the YP.24H variant (Fig. 1). According to the GO terms, the genes of this module are involved in photosynthesis, spermidine biosynthesis, nucleosome assembly, metal ion transport, and glycolytic processes (Fig. 2). Among the top 5 hub genes of M3 was the regulator of oxidative stress responses. Its expression increased in the WD variant only in the S29 variety, on the other hand it was significantly reduced in YP.6H, YP.24H and S29.6H variants (Fig. 3). Expression of the hub gene DnaJ was significantly reduced in both YP.6H and YP.24H variants, while for S29 variety it was reduced only in the S29.6H variant (Fig. 3). The nodes of maximum adjacency in PPI-network of M3 had a relatively low degree of centrality (15, Supplementary Data S2).

Module 4 (M4) consisted of 67 genes (Supplementary Data S1). In decreasing order of the number of M4 genes, homeologous groups of chromosomes were distributed as follows: 3 (14 genes) = 7 (14) > 2 (12) = 6 (12) > 1 (6) = 5 (6) > 4 (3). Under the WD conditions, S29 and YP varieties were similar in M4 gene expression pattern, but differed significantly in responses to low temperature treatment (Fig. 1). In S29, module M4 was activated by cold treatment for 6 hours, and especially by 24-hour treatment, while in YP this module was activated to much less extent. According to the GO terms, M4 genes are involved in metabolic processes related to glycine metabolism, cell wall biogenesis, and metal ion transport (Fig. 2). The expression of two homeologs of the cysteine protease inhibitor (hub genes) changed only in response to low temperature, and in S29 their expression was more activated in the 24H variant (Fig. 3). Expression of two serine hydroxymethyl transferase homeologs (hub genes) increased in response to low temperature only in YP. Expression of the BTB/POZ and TAZ domain protein (hub gene) was significantly activated under the 24H conditions only in S29 (Fig. 3). The degree of centrality of M4 hub genes in PPI network was the highest (1000, Supplementary Data S2).

Module 5 (M5) consisted of 53 genes (Supplementary Data S1). In decreasing order of the number of M5 genes, homeologous groups of chromosomes were distributed as follows: 3 (19 genes) > 4 (10) = 5 (10) > 6 (7) > 2 (3) > 1 (2) > 7 (1). In the 24H variant, M5 was more strongly activated in S29 compared to YP (Fig. 1). According to the GO terms, M4 genes are involved in photosynthesis, nucleosome assembly, and redox homeostasis regulation (Fig. 2). Among the top 5 hub genes of M5 were: two calmodulin homeologs with the EF-hand domain, UDP-glycosyltransferase, photosystem II protein Psb28, and a protein with unknown function. Expression of M5 hub genes did not change under the WD, but increased under the lowtemperature treatment. In the YP variety, hub genes expression increased more actively compared to S29 (Fig. 3). The centrality degree of hub genes in PPI network was relatively high (200, Supplementary Data S2).

Module 6 (M6) consisted of 40 genes (S1). By the number of M6 genes, homeologous chromosome groups were distributed as follows: 4 (8 genes) = 6 (8) > 5 (7) = 1 (7) > 2 (6) > 7 (4). The varieties differed in M6 gene expression in the WD variant; in S29, the expression of these genes increased, while in YP it did not. At low temperatures, M6 gene expression decreased in both varieties, but more so in S29. Of the top five M6 hub genes, four were light-harvesting complexes of the photosystems and one was the EF-hand calcium-binding domain. No protein-protein interactions were found for M6 proteins.

### Co-expressed genes associated with signaling and stress response

Analysis of the co-expressed DEGs associated with signaling and stress response revealed more intervarietal differences (Fig. 4). The varieties differed in the expression of EXORDIUM-like 1 proteins (M4), associated with sucrose signaling. Under the WD, the expression of two EXORDIUM-like 1 protein genes was suppressed in the S29 variety, but not in the YP. Under the 6H conditions, the expression of two EXORDIUM-like 1 protein genes was activated in YP, but not in S29, and under the 24H conditions, three EXORDIUM-like 1 protein genes were upregulated in YP.

**Fig. 4.**
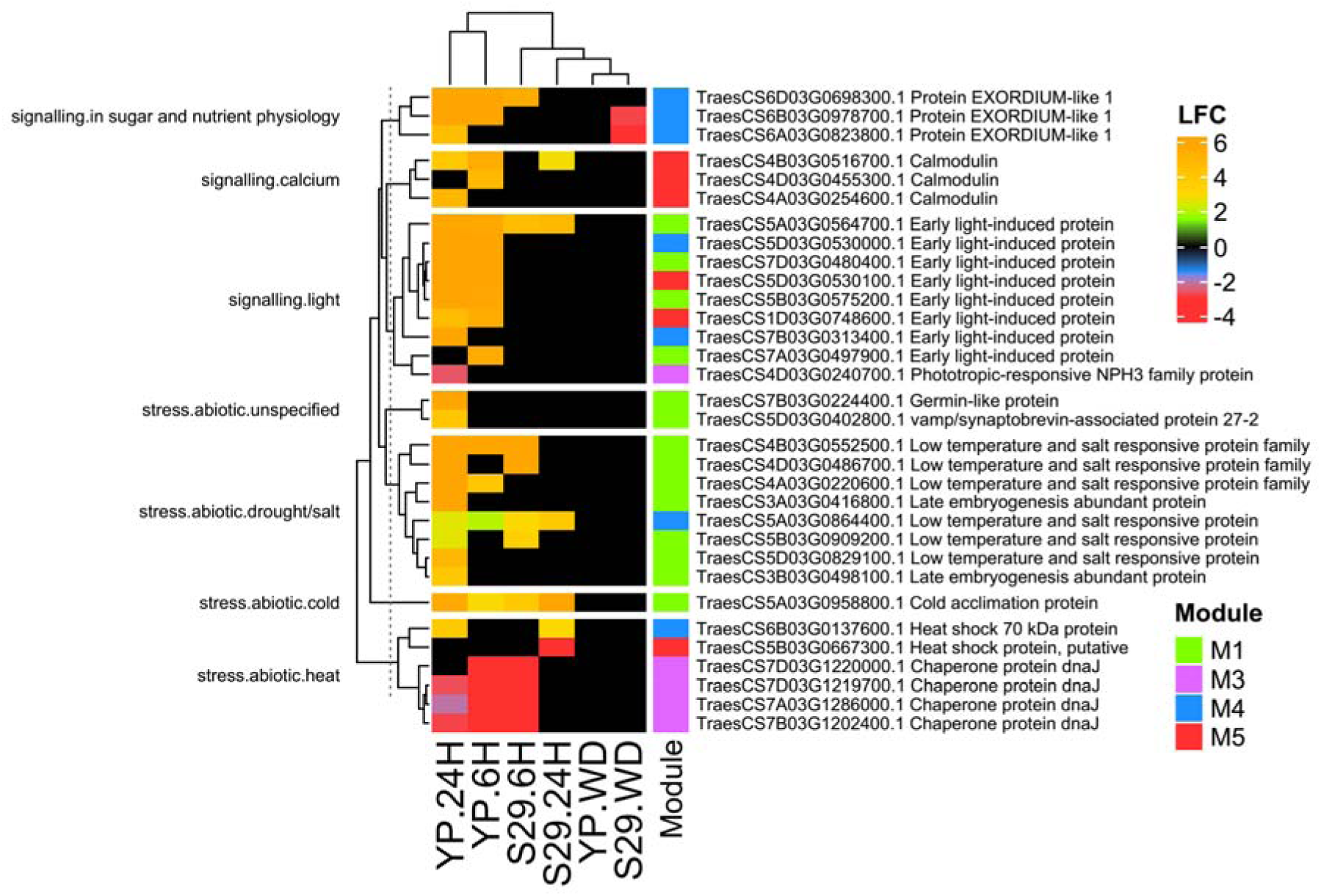
The expression of signaling and stress response-related co-expressed genes. LFC stands for Log Fold Change.

In YP compared to S29, more calmodulin (M5) genes involved in calcium signaling and genes associated with light signaling were upregulated under the low temperature treatment. Other groups of signaling-related genes also showed the greatest expression changes in YP in response to cold stress. S29 had fewer genes with DE, and their number decreased with increasing cold exposure, while in YP, conversely, it increased. The chaperone protein DnaJ was downregulated in both varieties under the 6H treatment, while under the 24H only the YP showed such expression pattern.

### The correlated gene expression network of S29 and YP wheat varieties

For the gene network analysis, we took datasets of co-expressed genes. Fig 5 shows correlated DEGs of wheat varieties S29 and YP under the WD (A, D), 6H (B, E) and 24H (C, F) conditions.

**Fig. 5.**
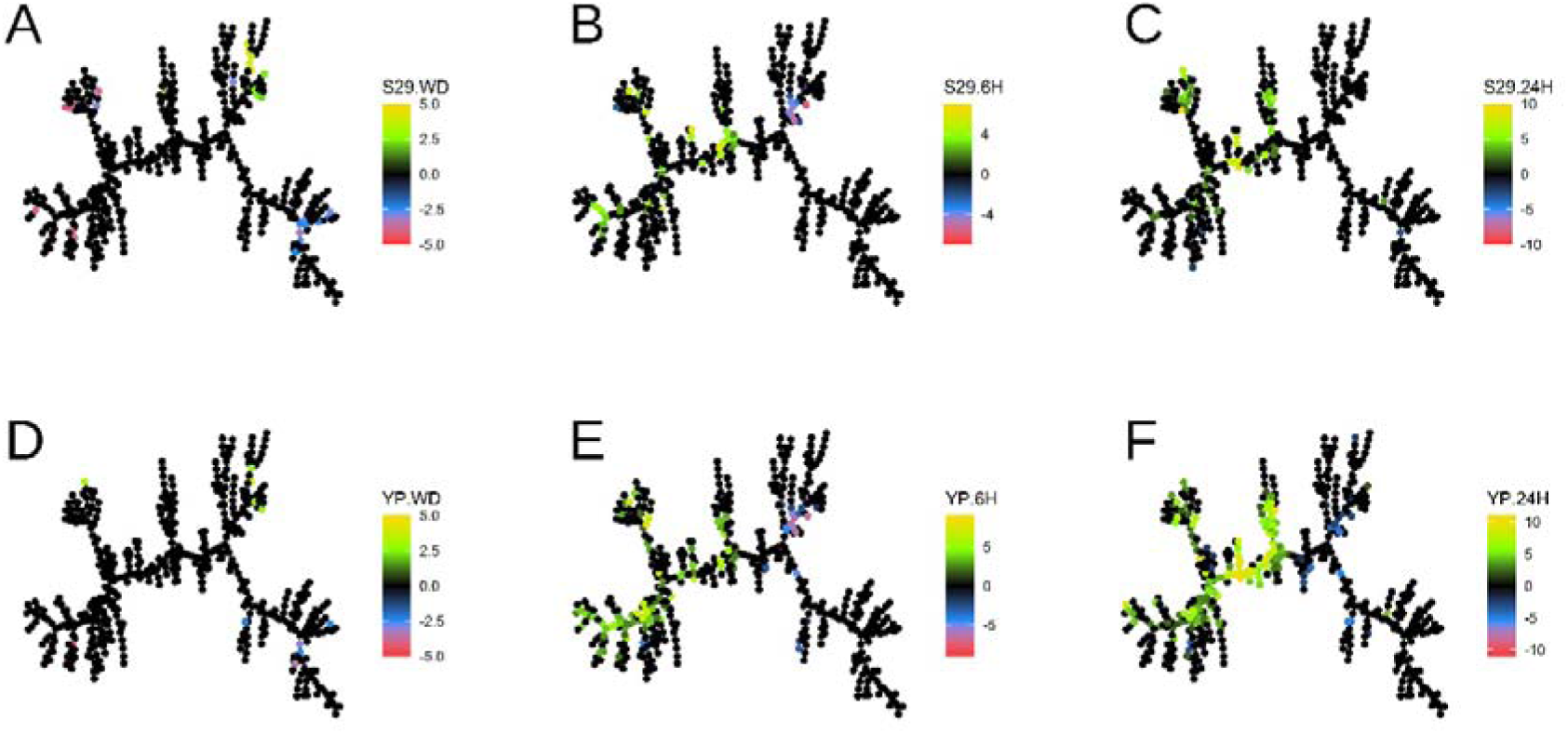
The minimum-spanning tree representation of correlated gene expression network with node colors according to Log Fold Change in wheat varieties S29 and YP under the WD (A, D), 6H (B, E) and 24H (C, F) conditions.

### Correlated gene expression in S29 and YP under the WD

The WD treatment induced more extensive transcriptomic changes in S29 than in YP (Fig. 5 A, D). Figure 6 shows two fragments of the gene network that showed the greatest differences between varieties in gene DE under the WD. In S29, four genes were upregulated under the WD, and the *TraesCS3D03G0906100* gene encoding the key enzyme of proline biosynthesis, delta1-pyrroline-5-carboxylate synthase, was upregulated in both S29 and YP varieties (Fig. 6 A).

**Fig. 6.**
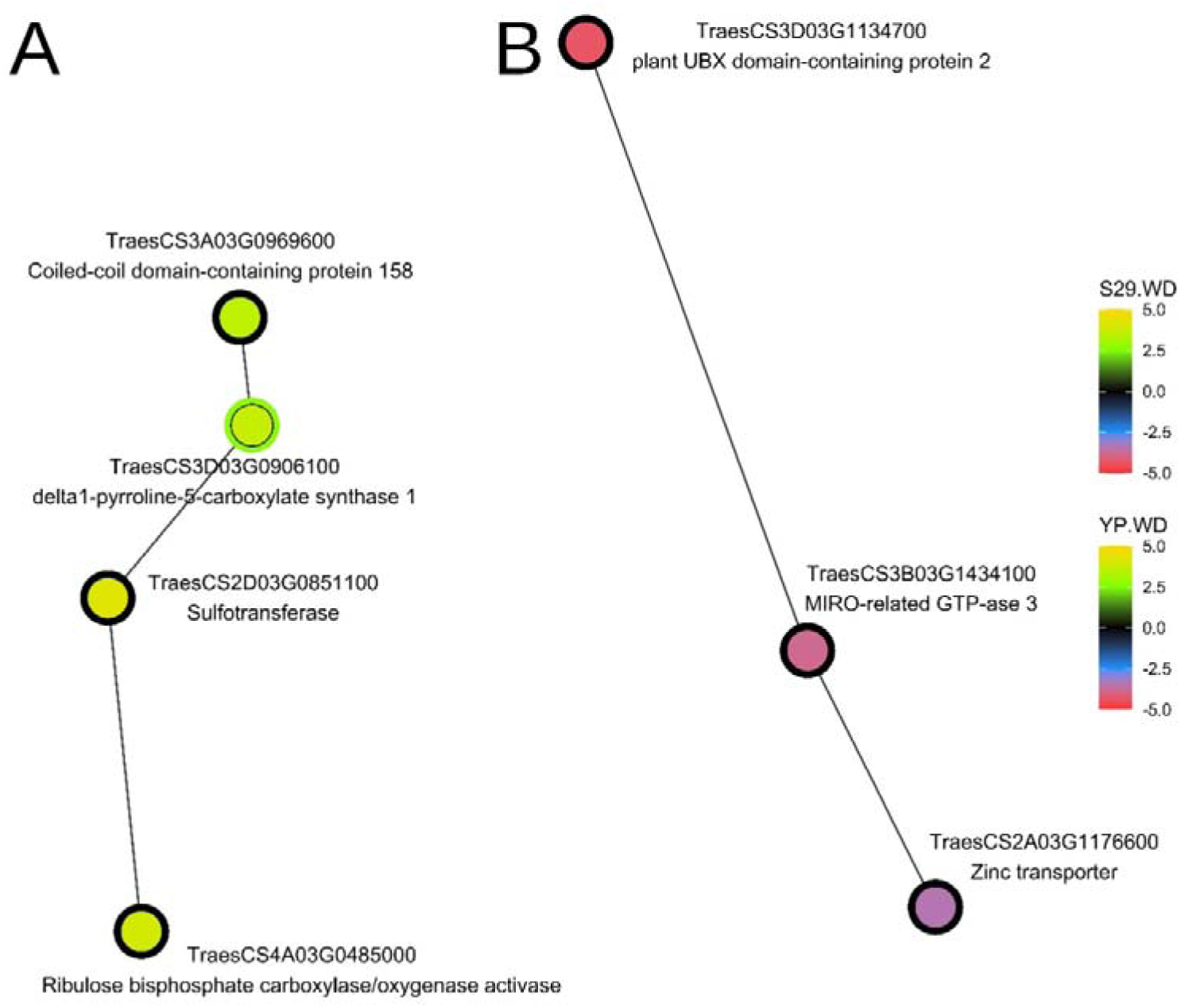
Fragments of gene network MST with genes (nodes) differentially expressed under the WD in YP and S29. The inner node color represents LFC in S29, the outer node color – in YP.

Three highly correlated genes *TraesCS2D03G0851100.1*, *TraesCS4A03G0485000.1* and *TraesCS3A03G0969600,* encoding sulfotransferase, Rubisco activase, and a protein with a Coiled-coil domain, respectively were co-upregulated with *TraesCS3D03G0906100* in S29, but not in YP. The greatest differences between varieties were observed in the expression of Rubisco activase (M3). In S29 compared to YP, three highly-correlated M4 genes were downregulated: *TraesCS3D03G1134700.1, TraesCS3B03G1434100.1,* and *TraesCS2A03G1176600.1*, encoding plant UBX domain-containing protein 2, MIRO-related GTP-ase 3, and zinc transporter, respectively (Fig. 6, B).

### Correlated gene expression in S29 and YP under the 6H

The transcriptional response to low positive temperature for 6 hours was stronger in the YP variety compared to S29 (Fig. 5 B, E). A fragment of the gene network (Fig. 7 A) shows differences between the varieties in expression of *TraesCS5A03G1103200* and *TraesCS5D03G1060100* encoding the NAC domain-containing proteins.

**Fig. 7.**
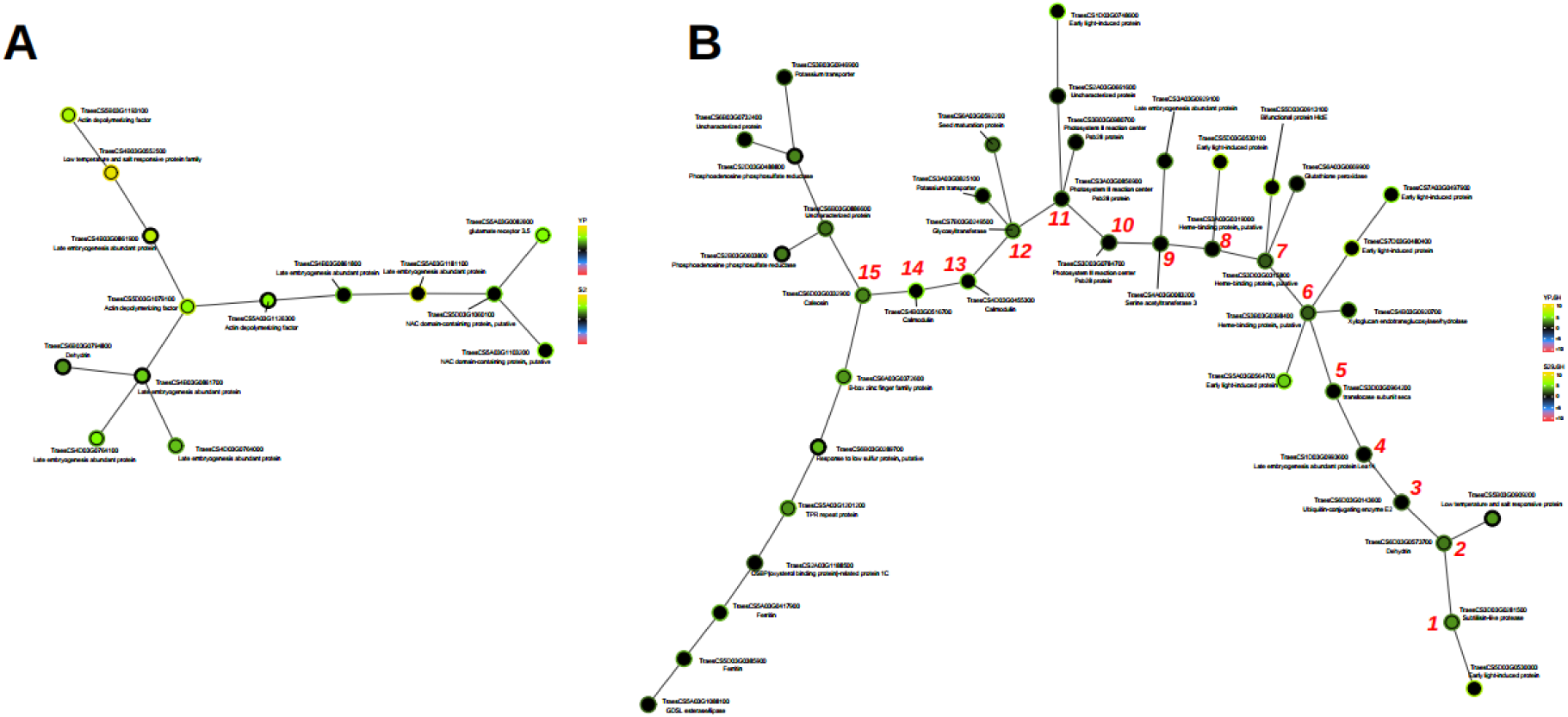
Fragments of gene network MST with genes (nodes) differentially expressed under the 6H in YP and S29. The inner node color represents LFC in S29, the outer node color – in YP.

Another intervarietal difference was noted in the expression of the M1 hub genes encoding actin depolymerizing factor. In S29, all three homeologs (*TraesCS5A03G1126300, TraesCS5B03G1193100, TraesCS5D03G1079100*) were upregulated, while in YP, only two of them (located in chromosomes 5B and 5D) were upregulated. Different late embryogenesis abundant (LEA) protein genes were activated in the varieties, but the total number of these genes was five in both of them.

In YP, a large backbone structure of highly-correlated genes consisting 15 nodes with branches was upregulated, whereas in S29, only nodes with numbers 1, 2, 6, 7, 12, and 15 of this structure were upregulated (Fig. 7, B). Unlike in S29, in YP the branches of nodes 1, 6, and 8, four genes (*TraesCS5D03G053100, TraesCS5D03G0530000, TraesCS7D03G0480400,* and *TraesCS7A03G049790*) encoding early light-induced proteins were upregulated. Another noted intervarietal difference were the module 4 gene *TraesCS5D03G0913100* (node 7 branch), the *TraesCS4B03G0455300* and *TraesCS4B03G0316700* genes (nodes 13 and 14), and the *TraesCS6B03G0289700* gene (node 15 branch), encoding the bifunctional protein HIdE, calmodulins, and response to low sulfur protein, respectively.

### Correlated gene expression in S29 and YP under the 24H

The transcriptomic response to 24 hours of low temperature was significantly more active in the YP variety compared to S29 (Fig. 5 C, F). In YP, a long backbone structure of 18 genes (nodes 1-18) was upregulated, whereas in S29, only 8 nodes of this structure were upregulated (Fig. 8 A).

**Fig. 8.**
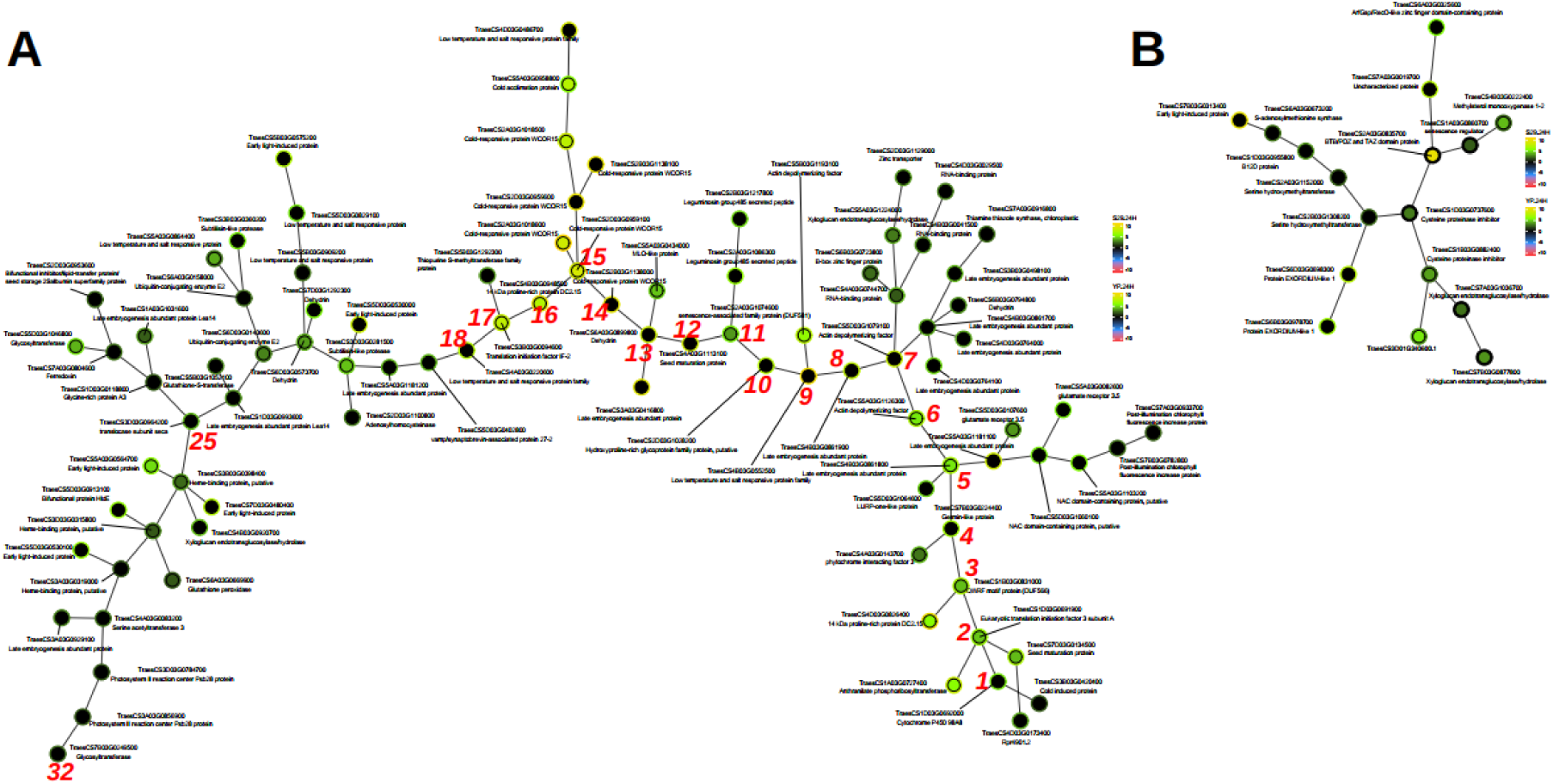
Fragments of gene network MST with genes (nodes) differentially expressed under the 24H in YP and S29. The inner node color represents LFC in S29, the outer node color – in YP.

In YP, more hub genes (encoding LEA protein, actin depolymerizing factor, and cold-responsive protein WCOR15) were activated. Unlike in S29, in YP, expression of two hub genes encoding the low temperature and salt responsive protein family was activated (Table 1).

**Table 1.** Genes of nodes 1–18 (Fig. 8, A), upregulated under the 24H in YP. Genes upregulated in both YP and S29 are highlighted in red.

| N | GeneID | Description |
| --- | --- | --- |
| 1 | TraesCS1D03G0692000 | Cytochrome P450 98A8 |
| 2 | **TraesCS1D03G0691900** | Eukaryotic translation initiation factor 3 subunit A |
| 3 | **TraesCS1B03G0831000** | QWRF motif protein (DUF566) |
| 4 | TraesCS7B03G0224400 | Germin-like protein |
| 5 | **TraesCS4B03G0861800** | Late embryogenesis abundant protein |
| 6 | **TraesCS5A03G1126300** | Actin depolymerizing factor |
| 7 | TraesCS5D03G1079100 | Actin depolymerizing factor |
| 8 | TraesCS4B03G0861900 | Late embryogenesis abundant protein |
| 9 | TraesCS4B03G0552500 | Low temperature and salt responsive protein family |
| 10 | TraesCS2D03G1028200 | Hydroxyproline-rich glycoprotein family protein, putative |
| 11 | **TraesCS2A03G1074600** | senescence-associated family protein (DUF581) |
| 12 | TraesCS4A03G1113100 | Seed maturation protein |
| 13 | TraesCS6A03G0899800 | Dehydrin |
| 14 | TraesCS2B03G1138000 | Cold-responsive protein WCOR15 |
| 15 | **TraesCS2D03G0959100** | Cold-responsive protein WCOR15 |
| 16 | **TraesCS4B03G0948500** | 14 kDa proline-rich protein DC2.15 |
| 17 | **TraesCS3B03G0094600** | Translation initiation factor IF-2 |
| 18 | TraesCS4A03G0220600 | Low temperature and salt responsive protein family |

More genes were also upregulated in YP node branches. Branches of the 5th node contained five module 1 genes, the expression of which was significantly more strongly activated in YP at 24H (Fig. 8A). Two of them (*TraesCS5A03G1103200* and *TraesCS5D03G1060100*) are orthologs of the AT5G63790 gene encoding NAC102 in Arabidopsis. Three genes with more active expression in YP (*TraesCS5A03G0082600*, *TraesCS5A03G1181100*, and *TraesCS5D03G1064600*) encode glutamate receptor 3.5, late embryogenesis abundant protein, and LURP-one-like protein, respectively. In addition, two node 11 branch genes (*TraesCS2A03G1086300* and *TraesCS2B03G1217800*) encoding leguminosin group 485 secreted peptide were more strongly activated in YP.

In YP all four nodes neighbor to node 15, encoding cold-responsive protein WCOR15, were upregulated, whether in S29 only two of them were (*TraesCS2A03G1018600 u TraesCS2D03G1018500*). More than that, neighbor node of low temperature and salt responsive proteins family (*TraesCS4D03G0486700*) was upregulated in YP.

The left part of a big backbone structure contains intervarietal expression difference in 11 genes (Fig. 8A), including genes encoding the light-induced protein (4), low temperature and salt responsive protein family (2), subtilisin-like protease, and other genes. In YP, strong upregulation of *TraesCS5D03G0913100* (bifunctional protein HIdE) occurred, just as it was noted for 6 hours if cold treatment.

A small fragment of the gene network demonstrates differences in the expression of M4 genes between varieties (Fig. 8, B). Dramatic differences were observed in the DE of the *TraesCS2A03G0835700* gene, encoding BTB/POZ and TAZ domain protein (S1). This gene was correlated with the expression of two M4 genes (*TraesCS6A03G0325600* and *TraesCS7A03G00019700*), encoding ArtGap/RecO-like zinc finger domain-containing protein and an uncharacterized protein, as well as the *TraesCS4B03G0222400* and *TraesCS1A03G0860700* genes, encoding methylsterol monooxygenase 1-2 and senescence regulator. In addition, S29 and YP showed significantly different DE of two M4 genes (*TraesCS7A03G1036700* and *TraesCS7B03G0877800*) encoding xyloglucan endotransglucosylase/hydrolase, two genes (*TraesCS6B03G0978700* and *TraesCS6D03G0698300*) encoding EXORDIUM-like 1 protein, as well as the M4 genes *TraesCS1D03G0737600* and *TraesCS7B01G120900*.1 encoding cysteine proteinase inhibitor and early light-induced protein. Overall, transcriptome reprogramming at low positive temperatures was more extensive and prolonged in YP compared to S29.

## Discussion

The bread wheat varieties used in our study were developed using conventional breeding for different geographic regions. The drought-tolerant, extensive variety S29 is approved for use in ten regions of the Russian Federation with high climatic risk. The intensive variety YP was developed for the temperate continental climate of eastern Germany. Compared to S29, YP has lower physical dough properties and is highly sensitive to abiotic stress factors (Pshenichnikova et al., 2012). The varieties differ significantly in quantitative characteristics of leaf pubescence. S29 has densely pubescent leaves, while YP has significantly fewer and shorter leaf trichomes (Doroshkov et al., 2016). According to long-term observations, the balance between the rate of CO_2_ assimilation (A) and the rate of transpiration (E) (A/E) in the S29 variety was stable under different water supply conditions, in contrast to the YP variety, in which this balance tended to decrease under drought conditions (Osipova et al., 2024).

Plant tolerance to drought and low temperatures is determined by complex, multilevel regulatory mechanisms involving perception, signaling, and stress response. These mechanisms are typically specific to drought and low temperature stress (Kim et al., 2024). Transcriptomic responses to abiotic stressors, which determine genotype tolerance to changes in external conditions, have also been shown to be specific for different bread wheat varieties (Amirbakhtiar et al., 2021; Chu et al., 2021; Iquebal et al., 2019; Niu et al., 2023; Prasad et al., 2022). Transcriptome analysis of S29 and YP cultivars at the seedling stage revealed 43 genes with conserved expression patterns in response to water deficit and low temperature. These include genes encoding the DnaJ chaperone protein, WRKY and bZip transcription factors, genes involved in cell wall biogenesis (glucan endo-1,3-beta-glucosidase and xyloglucan endotransglucosylase/hydrolase), genes encoding transmembrane proteins, cellular receptors, and others (Konstantinov et al., 2021). However, a relatively low overlap between the DEG lists was found, suggesting utilization of variety-specific regulatory mechanisms under different types of stress conditions. Using gene regulatory network analysis, we found differences in expression of highly-correlated co-expressed genes under water deficit and low temperature in extensive (S29) and intensive (YP) wheat varieties at the early development stage (Fig. 5).

### Differences in transcriptome responses of S29 and YP varieties to water deficit

Under the water deficit conditions, both varieties demonstrated relatively low DEG numbers in leaf tissue. Thirty genes were differentially expressed in leaves of variety S29, 12 of which were located in chromosomes of the second homeologous group. In the YP variety, only 11 genes were differentially expressed, and these were evenly distributed across the chromosomes of the six homeologous groups, excluding the first.

Two fragments of the gene network under the WD demonstrate the most significant intervarietal expression differences of several genes assigned to co-expression modules 3 and 4 (Fig. 6 A, B). In contrast to YP, in S29 the expression of M4 genes encoding plant UBX domain-containing protein 2 and Miro-related GTPase 3 was downregulated. Proteins containing the UBX domain are known to be involved in the temporal and spatial regulation of the activity of the AAA-ATPase Cdc48/p97 (Schuberth & Buchberger, 2008). The chaperone-like complex Cdc48/p97 is required for numerous processes of ubiquitin-dependent degradation of proteins associated with endoplasmic reticulum, mitochondria, and nucleus (Li et al., 2022). GTPases of the Miro family are known to be involved in establishing and maintaining contact sites between mitochondria and the endoplasmic reticulum (ER), which plays a crucial physiological role in a variety of mitochondrial processes, including mitochondrial transport and distribution of mitochondria within the cell (Kanfer & Kornmann, 2016; Michel & Kornmann, 2012). They also play a key role in signaling pathways, control mitochondrial dynamics and cytoskeletal maintenance (Aspenström, 2024; Zinsmaier, 2021), and participate cellular stress response process, which is undoubtedly important for plant adaptation to changing conditions. Suppression of UBX2 and Miro-related GTPase 3 expression indicates that S29 cells under the WD saved a significant amount of ATP and GTP energy by reducing the activity of energy-intensive processes such as (i) ubiquitin-dependent protein degradation in organelles and (ii) regulation of multiple mitochondrial processes carried out with the participation of Miro-related GTPases (Aspenström, 2024; Lee et al., 2016; Zinsmaier, 2021).

Among the three genes of M3, upregulated only in the S29 cultivar (Fig. 6A), was the Rubisco activase gene (*RCA*). This enzyme uses the energy of ATP hydrolysis to restructure Rubisco and restore its catalytic activity (Perdomo et al., 2024). Upregulation of *RCA* in S29 apparently maintained Rubisco activity in the face of decreased expression of the small subunit structural genes (Fig. 3) and, consequently, a decrease in the native enzyme content. These data indicate a redistribution of ATP energy consumption in S29 cells to maintain the activity of Rubisco, the key enzyme of the Calvin–Benson cycle responsible for CO_2_ assimilation. No similar activation of *RCA* expression occurred in the leaves of the YP variety.

More efficient regulation of the Calvin–Benson cycle in S29 under the WD conditions was also indicated by the suppression of chloroplast protein CP12 expression, with the suppression of the homeolog from chromosome 2A (Supplementary Data S1) being particularly significant. CP12 is a linker protein that binds glyceraldehyde-3-phosphate dehydrogenase (GAPDH) and phosphoribulokinase (PRK) into a single complex in which the activity of GAPDH and PRK is inhibited (Gérard et al., 2022). Suppression of CP12 expression in S29 under the WD contributed to the maintenance of an unlinked, active state of GAPDH and PRK, which apparently created more favorable conditions for the functioning of the Calvin–Benson cycle under the WD.

The expression of proteins with the coiled-coil (CC) domain and a sulfotransferase was found to be highly-correlated with *RCA* expression. CC-domain proteins are known to be involved in immune signaling in plants and can interact with transcription factors of the WRKY, MYB, and Golden2 (GLK) families (Townsend et al., 2018; Wang et al., 2021). The correlated activation of this gene’s expression in S29 could indicate its involvement in the transcriptional regulation of Rubisco activase and sulfotransferase, an enzyme that plays a vital role in wheat plant growth and development and in responses to water stress (Chaudhary et al., 2023).

The downregulation of two genes encoding the EXORDIUM-like 1 protein, which has been shown in *Arabidopsis* to be part of a regulatory pathway that controls growth and development under carbon and energy starvation by Schröder et al. (Schröder et al., 2009; Schröder et al., 2011), in S29 indicates efficient coordination of growth and metabolism under drought stress in this variety.

Thus, the differences observed between the S29 and YP varieties in the transcriptome response to WD suggest that, unlike in YP variety, energy resources in leaves of the extensive variety S29 are redistributed to support the activity of the Calvin-Benson cycle, which is the primary pathway for carbon fixation in plants. We believe that these metabolic changes play a primary role in shaping the adaptive stress-response of the S29 variety.

### Differences in transcriptomic responses of S29 and YP varieties to low positive temperature treatment

The most pronounced intervarietal difference in DEG numbers was observed under prolonged exposure to low temperature (24H). In the S29 variety, 49 genes were differentially expressed, the largest number (10) of which were from chromosomes of the fifth homeologous group. In YP, 145 genes were differentially expressed under the same conditions. The largest number of genes (27 and 36) were expressed from chromosomes of the second and fifth homeologous groups, which is in a good agreement with the data of meta-analyses by Acuña-Galindo et al. (Acuña-Galindo et al., 2015) and Bhanbhro (Bhanbhro et al., 2024) on the localization of a large number of metaQTLs associated with physiological resistance traits in bread wheat on these chromosomes.

The studied varieties differed in the activation of several transcriptional regulators under low temperatures. Unlike S29, in YP variety genes encoding NAC-domain transcription factors from chromosomes 5A and 5D were upregulated both in response to 6H and 24H treatment. Unlike YP, in S29 the expression of M4 hub gene BTB/POZ and TAZ domain was upregulated at 24H. Activation of this gene expression from chromosome 2A explained a significant proportion of the intervarietal differences observed in module M4 at 24H. BTB/POZ is a versatile, highly interactive, and multifunctional protein that, in combination with the TAZ domain, is involved in plant growth and development as a transcriptional regulator (Shalmani et al., 2021). In recent years, its crucial role in the stress tolerance of sugar beet, rice, potato, and soybean plants has been demonstrated (Elsanosi et al., 2024; Feng et al., 2024; Mandal et al., 2023). The potential usefulness of the BTB/POZ genes for improving the responses of crop plants to multiple abiotic stressors has been noted (Mandal et al., 2023). Activation of the expression of this protein was specific to the S29 variety and indicated its important role in the adaptation of extensive wheat varieties to low temperatures. The expression of a serine protease inhibitor from chromosomes 1B and 1D was correlated with the activation of BTB/POZ and TAZ expression. More active expression of this gene in S29 indicated a balanced consumption of protein reserves by inhibiting proteolytic processes. As under the WD conditions, under low temperatures the extensive type S29 variety implemented a resource-saving strategy.

Under the 24H treatment, the varieties studied differed significantly in expression pattern of defense response genes (Figs. 5, 8). Obvious differences were found in the expression of early light-induced protein (ELIP) genes from chromosomes of homeologous groups 5 and 7, as well as from chromosome 1D. The upregulation of these genes occurred only in variety YP. The function of ELIPs is associated with protection from photooxidation of light-harvesting complexes in *Arabidopsis* chloroplasts (Hutin et al., 2003). In wheat plants, a sharp increase in ELIP transcript levels was shown at 4°C under low light conditions, and even in the dark (Shimosaka et al., 1999). The accumulation of these proteins at 24H in variety YP apparently served as a mechanism for protecting photosystems under chilling stress. In addition, a larger number of late embryogenesis abundant protein (LEA) genes were upregulated in the YP variety at 24H compared to S29. The best-studied function of LEA proteins is to protect the stress-responsive proteome (Dirk et al., 2020). In bread wheat, 281 LEA genes were identified to be regulated by abiotic stresses, especially salt and cold stress (Zan et al., 2020). Additionally, in YP variety more cold responsive protein WCOR15 genes, encoding a small protein targeting epidermal guard cell chloroplasts (Takumi et al., 2003), were found to be upregulated.

Based on differential expression of the serine hydroxymethyl transferase and S-adenosylmethionine transferase genes, the YP cultivar activated the synthesis of methyl donors for the formation of key metabolites and protein methylation. The YP cultivar also showed increased expression of ubiquitin-conjugating E2 proteins, whose involvement in responses to abiotic stress factors has been demonstrated in various plants (Liu et al., 2020), and glycine-rich cell wall structural proteins, which are involved in the development of cold tolerance by participating in the transduction of external signals and also exhibiting chaperonin activity toward RNA (Czolpinska & Rurek, 2018). In YP, at 24H, more M1 hub genes encoding actin depolymerizing factor (ADF), which remodulates the actin cytoskeleton and is involved in many cellular processes, including cold adaptation (Sun et al., 2023), and other genes were activated.

Important information on the genes upregulated in YP and S29 varieties under chilling stress is presented in Table 1. First and foremost, we found differences between the YP and S29 varieties in the number of upregulated at 24H genes in the gene network (18 in YP and 8 in S29). We believe that the increased expression of these eight genes in the extensive S29 variety and the intensive YP variety indicates universal mechanisms of response to chilling stress in different wheat varieties. This stress-response pathway appears to be promising for use in breeding programs to improve polygenic traits for resistance to abiotic stress in wheat. Overall, such hypothesis is supported by the existing studies on the role of these genes in resistance to abiotic and biotic stresses (Berna & Bernier, 1999; Chana et al., 2025; Koubaa & Brini, 2020; Kreps et al., 2002; Li et al., 2025; Morant et al., 2002; Singh et al., 2013; Takumi et al., 2003; Wang et al., 2019).

The relative expression levels of most of the genes discussed were higher in S29 variety than in YP under control conditions. Transcriptomic data indicate that the extensive variety S29 possessed constitutive molecular tolerance to both drought and low positive temperatures, while the high-yielding, intensive variety YP, sensitive to growing conditions, activated an expanded arsenal of molecular mechanisms to withstand cold stress.

In the future, it will be desirable to study the edaphic adaptation of the S29 and YP varieties, since it remains unknown which adaptive strategies are used by these wheat varieties at the transcriptome level under conditions of the full life cycle, which depends on soil conditions and occurs in continuous interaction of the plant organism with the soil microbiome.

## Conclusions

In this study, we identified for the first time the differences in molecular mechanisms of adaptation to drought and low positive temperatures in extensive (S29) and intensive (YP) wheat varieties. A distinctive feature was the DEGs of several regulatory proteins. In the S29 variety, drought stress activated the expression of proteins with the CC domain, while prolonged exposure to low temperature activated the expression of BTB/POZ and TAZ domain proteins. In the YP variety, low temperature activated transcription factors of the NAC family. Overall, the transcriptome response of the intensive YP variety to prolonged exposure to low temperature was more active and extensive compared to S29. The greatest contrast was observed in the DEGs of co-expression module M1, associated with stress response, and module M4, in which differences between varieties were determined mainly by the expression of BTB/POZ and TAZ domain proteins. A distinctive feature of the extensive S29 variety was its transition to energy-saving mode to maintain the activity of the Calvin-Benson cycle at low temperatures and a reduction in proteolytic processes at low temperatures. Understanding the diversity in the molecular mechanisms of adaptation to environmental stress factors in wheat varieties intended for different types of agriculture is essential for the development of marker-assisted and genomic wheat breeding technologies.

## Author Contributions

All authors contributed to the study conception and design. Data collection and analysis were performed by Igor Vladimirovich Gorbenko. The first draft of the manuscript was written by Svetlana Vladimirovna Osipova and all authors commented on previous versions of the manuscript. All authors read and approved the final manuscript.

## Data availability

The datasets generated during and/or analyzed during the current study are available in the github repository, https://github.com/ivg-git/IVG-variability-of-wheat-transcriptional-response-public-data.

